# When should bees be flower constant? An agent-based model highlights the importance of social information and foraging conditions

**DOI:** 10.1101/2022.07.02.498534

**Authors:** Lucy Hayes, Christoph Grüter

**Affiliations:** School of Biological Sciences, University of Bristol, 24 Tyndall Avenue, BS8 1TQ Bristol, UK

## Abstract

1. Many bee species show flower constancy, *i*.*e*. a tendency to visit flowers of one type during a foraging trip. Flower constancy is important for plant reproduction, but whether bees also benefit from flower constancy remains unclear. Social bees, which often use communication about food sources, show particularly strong flower constancy.
2. We hypothesised that the sharing of social information increases the benefits of flower constancy because foragers share information selectively about high-quality food sources, thereby reducing the need to sample alternatives. We also asked if foraging landscapes affect flower constancy. We developed an agent-based model that allowed us to simulate bee colonies with and without communication and flower constancy in different foraging environments.
3. Flower constancy alone performed poorly in all environments, while indiscriminate flower choice was often the most successful strategy. However, communication improved the performance of flower constant colonies in nearly all tested environments. This combination was particularly successful when high-quality food sources were abundant and competition was weak.
4. Our findings help explain why social bees tend to be more flower constant than solitary bees and suggest that flower constancy can be an adaptive strategy in social bees. Simulations suggest that anthropogenic changes of foraging landscapes will have different effects on the foraging performance of bees that vary in flower constancy.

## Introduction

Most flowering plant species are animal pollinated and bees, in particular, are important pollinators of wild and agricultural plants (Bawa, 1990; Klein et al., 2007; Ollerton et al., 2011). Several biological features explain why bees are helpful agents of reproduction for plants, including their abundance and their often broad (*i*.*e*. polylectic) diet in combination with a tendency to specialise on a particular flower type during an individual foraging bout. The latter behaviour, called flower constancy (Bateman, 1951; Chittka et al., 1999; Darwin, 1876; Grüter & Ratnieks, 2011; Waser, 1986), reduces conspecific pollen loss and heterospecific pollen deposition, both of which can reduce plant fitness (Ashman & Arceo-Gómez, 2013; Campbell & Motten, 1985; Chittka et al., 1999; Morales & Traveset, 2008; Waser, 1986). Flower constancy is also thought to enhance the coexistence of different plant species and, thus, shapes plant community structure (Morales & Traveset, 2008; Song & Feldman, 2014).

From a pollinator perspective, however, the benefits of flower constancy are less obvious. Ignoring potentially superior flower species appears to contradict optimal foraging theory (King & Marshall, 2022; Latty & Trueblood, 2020; Waser, 1986; Wells & Wells, 1983, 1986). Why then are pollinators flower constant? The most widely accepted view is that flower constancy is driven by cognitive limitations, which can include (*i*) slow learning to forage efficiently on a new flower species, (*ii*) an inability to memorise more than one or a few flower types, (*iii*) unstable short-term memories which are prone to being erased by competing information or (*iv*) an inability to retrieve long-term memory about different flower species fast enough to be an efficient generalist (Darwin, 1876; Heinrich, 1979; Lewis, 1986; Menzel, 1999; Raine & Chittka, 2007; Waser, 1986; for reviews see Chittka et al., 1999; Grüter & Ratnieks, 2011). These cognitive limitations are likely to cause time delays as a bee tries to extract nectar from a flower after switching from a different species and they may increase switching times (Chittka et al., 1999; Goulson et al., 1997; Lewis, 1986; Raine & Chittka, 2007).

The “cognitive limitations hypothesis” as an explanation for flower constancy is not without challenges. Given that efficient foraging is likely to be under strong natural selection due to its effects on reproductive success (Heinrich, 1979), why does natural selection not lead to the evolution of lower flower constancy in all bees? How can we explain the finding that individual bees often show plasticity in their flower constancy, e.g. by being more flower constant after finding good rewards (Chittka et al., 1997; Grüter et al., 2011; Wells & Rathore, 1994; but see Hill et al., 1997) or the distances between food sources are shorter (Gegear & Thomson, 2004; Kunin, 1993; Mardern & Waddington, 1981)? Why do bee species vary in their degree of flower constancy? Social bees, in particular, are often highly flower constant (Cholis et al., 2020; Free, 1963; Heinrich, 1976, 1979; Hill et al., 1997; Kozuharova, 2018; Pangestika et al., 2017; Rossi et al., 2015; Slaa et al., 2003; White et al., 2001; but see Martínez-Bauer et al., 2021), while flower constancy seems to be less pronounced in solitary bees (Bateman, 1951; Campbell & Motten, 1985; Eckhardt et al., 2014; Jakobsson et al., 2008; Pohl et al., 2011; Smith et al., 2019; Waser, 1986; Williams & Tepedino, 2003). Smith et al. (2019), for example, studied pollen composition of 56 bee species and found that individual social bees showed a higher degree of specialisation during a foraging bout than solitary species. Different ecological needs could explain this difference. Solitary bees need to collect all required nutrients by themselves, potentially favouring a strategy of mixing resources during a foraging trip even if this has energetic costs (e.g. Eckhardt et al., 2014; Williams & Tepedino, 2003). In social species, on the other hand, different bees from the same colony can specialise on different types of resources and flower species to cover their nutritional needs.

Foragers of many social bees share information about profitable food sources, and this could affect the value of flower constancy. Honeybees use the waggle dance to indicate the odour (type) and location of profitable food sources (von Frisch, 1967) and some stingless bees lay pheromone trails (Grüter, 2020; Jarau, 2009; Lindauer & Kerr, 1960; Nieh, 2004). Stingless bees and bumblebees inform nestmates about the availability and odour of a good food source by means of excitatory or jostling runs inside their nest (Dornhaus & Chittka, 2004; Hrncir, 2009). Trophallaxis – food transfer between bees – is performed by honey bees and stingless bees (Farina et al., 2012; Farina & Grüter, 2009; Hart & Ratnieks, 2002; Hrncir et al., 2006; Krausa et al., 2017; von Frisch, 1967) and is another behaviour that allows nestmates to learn the odour of available food sources (Aguilar et al., 2005; Farina et al., 2005; Lindauer & Kerr, 1960; von Frisch, 1967). These different communication behaviours share two common features. First, they depend on food quality. Dances, pheromone trails, jostling runs and trophallaxes occur at higher frequencies if the exploited food sources are of higher quality (Farina et al., 2012; Hrncir, 2009; Krausa et al., 2017; Lindauer & Kerr, 1960; von Frisch, 1967). Second, during these social interactions, nestmates can learn the odour of the exploited flower species and acquire a preference for this flower species in the field.

Heinrich (1976) was probably the first to propose a link between recruitment communication and flower constancy. Since recruiting bees share information selectively about high-quality food sources, recruits can discover profitable flower types without the costs of sampling different, lower-quality plant species. This does not require that foragers are able to direct nestmates to a specific location, as in honeybees and some stingless bees, but depends more generally on foragers biasing the food search towards flower types that are more profitable. Among the social bees, bumble bees seem to be less flower constant that honey bees (Bateman, 1951; Grant, 1950; Martínez-Bauer et al., 2021; Smith et al., 2019), possibly because their communication system is less sophisticated than that of honey bees (Heinrich, 1976).

Experimental studies of the benefits of flower constancy and how they depend on social and ecological traits are logistically challenging for several reasons. For example, it is often not possible to manipulate the degree of flower constancy while keeping other factors constant. Agent-based simulation models can be a useful complementary tool to evaluate how biological and ecological factors affect the benefits of a behavioural strategy. We developed an agent-based simulation model to test the hypothesis that flower constancy is more beneficial in bees that communicate about profitable food sources than in bees without communication. Colonies consisting of virtual bees (agents) were either flower constant or they chose food sources randomly (indiscriminately) in environments that varied in the number and quality of food sources. Some studies have found that bees adjust the degree of flower constancy depending on the foraging conditions, being more flower constant if the rewards on offer are better (Chittka et al., 1997; Grüter et al., 2011; Wells & Rathore, 1994; but see Hill et al., 1997) and the distances between food sources (or density) are shorter (Gegear & Thomson, 2004; Kunin, 1993; Marden & Waddington, 1981). We, therefore, expected flower constancy to be more beneficial in environments with more flowers and larger reward sizes. Exploitation competition, on the other hand, is expected to favour a indiscriminate choice because greater competition increases the costs of rejecting a reward (Pulliam, 1974).

## The model

We built an agent-based model (ABM) using the programming software NetLogo 6.1 (Wilensky, 1999) (see supplementary material for NetLogo file). It is an extension and further analysis of the model developed in Grüter & Hayes (in preparation), which analysed foraging distances. The model simulates an environment with a colony surrounded by food sources. The agents (“bees”) operate on a two-dimensional square grid with 400 × 400 patches. A single patch length corresponds to 5 meters and 1 tick corresponds to 1 second. Thus, the size of the virtual world corresponds to 2 × 2 km. The nest of the colony is positioned in the centre of the grid (x=0, y=0). In the default situation, environments contained two different flower types that differed in the rewards they offered.

The model allows simulating a wide range of parameter values, but for the purpose of this study we based our default parameters, such as the nest stay time (*t*_nest_), flight speed (*v*_flight_), metabolic costs of flying (*M*_cost_), and crop capacity (*Crop*) on the Western honeybee *Apis mellifera* because we have accurate information about these relevant biological parameters in *Apis mellifera*. Other values were tested (see Table 1 and section *Sensitivity analysis and model exploration*).

**Table 1:** Overview of the model variables and the used values.

| Variables | Description | Default values | Other values tested | Information source |
| --- | --- | --- | --- | --- |
| Colony size | Number of foragers in a colony | 100 | 5-300 | Westphal et al. 2006; Grüter 2020 |
| $FS_{\text{number}}$ | Number of food sources per type | 3000 | 1500, 4500 | arbitrary |
| $FS_{\text{size}}$ | Size of food sources | 1 patch | | arbitrary |
| $FS_{\text{types}}$ | Number of food source types | 2 | 4 | arbitrary |
| $Reward_{\text{HQ}}$ | Nectar amount per food source | 5μl | 2.5μl, 10μl | Willmer 2011 |
| $Reward_{\text{LQ}}$ | Nectar amount per food source | 2.5μl | 1.25μl, 5μl | Willmer 2011 |
| $v_{\text{flight}}$ | Flight speed | 1.4 patch/tick | | von Frisch 1967 |
| $M_{\text{cost}}$ | Metabolic costs of flight, J/tick | 0.032 | 0.016, 0.064 | Heinrich 1975; Willmer 2011 |
| $t_{\text{flower-stay}}$ | Time spent at food source | 60 ticks | 20, 180 | arbitrary |
| $v_{\text{nest}}$ | Movement speed inside nest | 0.1 patch/tick | | arbitrary |
| $t_{\text{nest-stay}}$ | Time in nest between trips | 300 ticks | 150, 450 | Farina 2000 |
| $Crop_{\text{HQ}}$ | Crop load when flower constant to HQ food sources | 50μl | 25 μl, 100 μl | Núñez 1966 |
| $Crop_{\text{LQ}}$ | Crop load when flower constant to LQ food sources | 25μl | 12.5 μl, 50μl | Núñez 1966 |
| $Crop_{\text{Random}}$ | Crop load without flower constancy | 37.5μl | 50μl | Núñez 1966 |
| $t_{\text{refill}}$ | Time until food sources offer food again | 0 ticks | 1200, 3600 | Stout & Goulson, 2002 |
| Lévy $\mu$ | Lévy flight parameter | 2.4 | 1.8, 3.0 | Reynolds et al. 2009 |
| $Constancy_{\text{HQ}}$ | Probability to remain constant to a type after leaving a high-quality food source during flower constant simulations | 100% | | Grüter et al. 2011 |
| $Constancy_{\text{LQ}}$ | Probability to remain constant to a type after leaving a low-quality food source during flower constant simulations | 100% | 90%, 95% | Grüter et al. 2011 |

Foragers in social bees use different behavioural mechanisms to transmit social information and, thereby, influence the food source preferences of their nestmates (see introduction). The model does not simulate a particular behaviour, but a generic process that biases the food preferences of nestmates, which could correspond to jostling runs, trophallaxis or the waggle dance.

### Entities and state variables

#### Bees

The default colony size was 100 agents (forager bees), which corresponds to the size of the forager pool in many species of bumble bees (Westphal et al., 2006) and stingless bees (Grüter, 2020). Agents could assume any of the following states: (1) *generalists*, (2) *feeding forager, (3)searching forager*, (4) *returning forager*, (5) *inside-nest-worker* and (6) *influencer* (see Fig. 1).

**Fig. 1.**
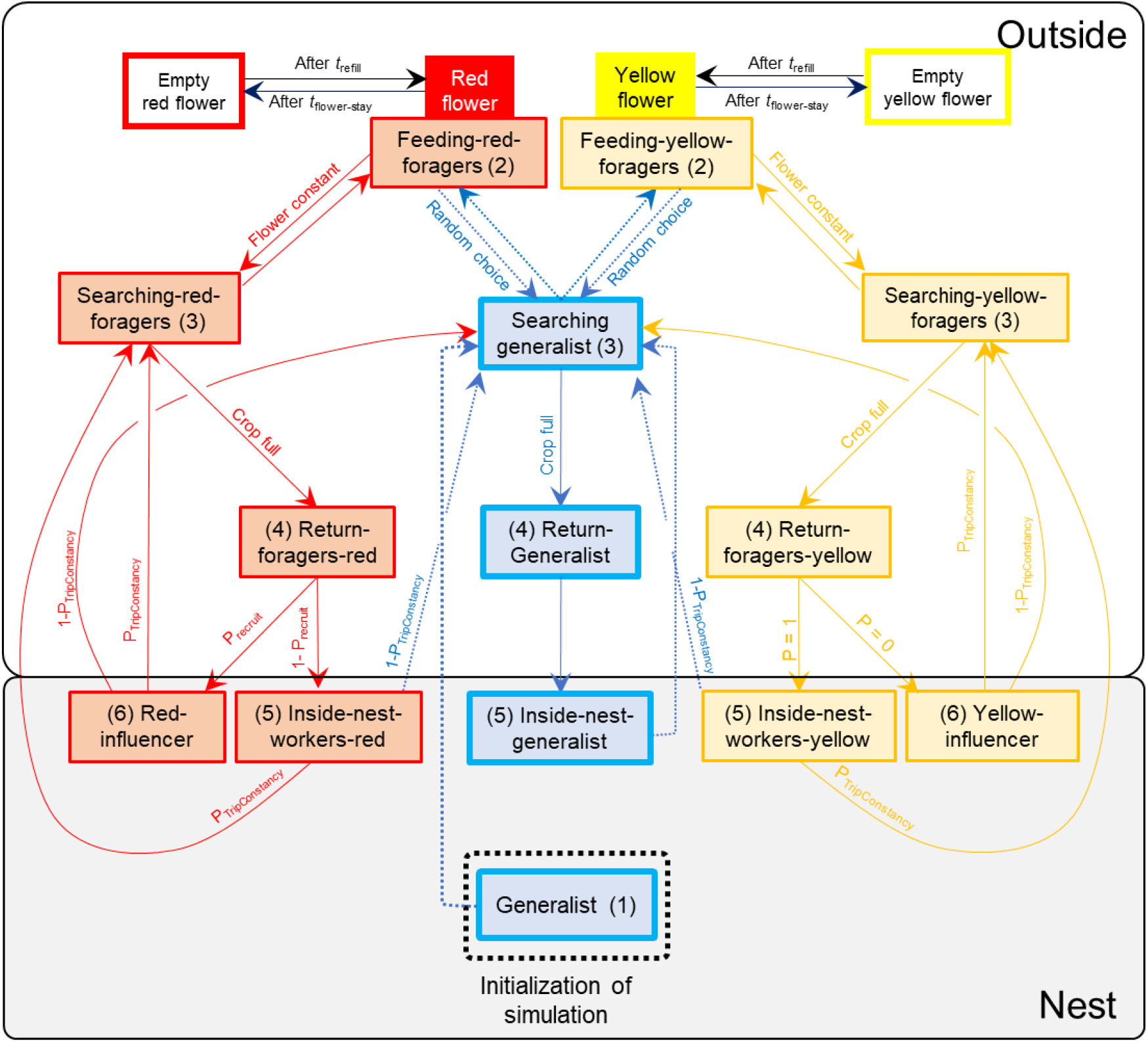
State diagram showing the different states of the agents and the possible transitions between states. Here, yellow flowers were arbitrarily chosen to represent a lower-quality food source, therefore, the default probability that foragers visiting yellow flowers would become influencers after their return to the nest was 0.

Agents begin the simulation in the centre of the nest with energy = 0 as *generalists*. They then move at a flying speed of 1.4 patch/tick (*v*_flight_), corresponding to a flight speed of *Apis mellifera* (7 m/sec, von Frisch 1967). Their random search behaviour follows a Lévy-flight pattern (with *μ* = 2.4 as default) (Reynolds, 2009; Reynolds et al., 2007). A Lévy-flight consists of a random sequence of flight segments whose lengths, *l*, come from a probability distribution function having a power-law tail, *P*(*l*)∼*l*^−μ^, with 1<μ<3 (Reynolds et al., 2007). The speed of agents moving inside the nest (*v*_nest_) was arbitrarily chosen to be 0.1 (patch/tick). Flying has a metabolic cost (*M*_cost_) of 0.032 Joule (J) per tick in the default condition (Heinrich, 1975; Willmer, 2011). Once an agent encounters a food source, they remain on the food source for 60 ticks (*t*_flower-stay_) under default conditions (*feeding foragers*), irrespective of whether they were choosing indiscriminately or are flower constant. Thus, we assume that the time spent handling a flower or flowers in a patch and extracting the reward is the same for flower constant and indiscriminate foragers. This was chosen as the default condition to explore whether flower constancy can be an adaptive strategy in the absence of cognitive constraints.

The agent then continues to forage (*searching foragers*) until its crop is full, after which it returns to the nest (*returning foragers*) to unload its energy and stay in the nest for 300 ticks (*t*_nest-stay_) (Farina, 2000; Seeley, 1986; von Frisch, 1967). In the default condition, only foragers visiting the high-quality food source could become *influencers* (*i*.*e*. bees that bias the food choice of other bees) upon return to the nest. *Influencers* target *inside-nest-workers* that are not yet flower-constant to the high-quality food type by changing the latter’s preference if they encountered each other on same patch inside the nest. Following such an encounter, *inside-nest-workers* become flower constancy for the high-quality type.

Since recruitment behaviours often depend on the food source distance (with greater foraging distances lowering the probability of recruitment), we simulated recruitment curves where the probability of becoming an *influencer* decreased with increasing distance of the last visited food patch (Fig. S1).

### Food sources

In the default condition, two different types of food sources can be found in the environment, mimicking the typical situation in experimental flower constancy studies (e.g. Chittka et al., 1997; Goulson & Wright, 1998; Grüter et al., 2011; Ishii & Masuda, 2014; Wells & Wells, 1983). The food source types differ in the rewards they offer per visit. Natural bee-visited flowers offer between 0.1 and 10 μL of nectar per flower (Willmer, 2011, p. 203). For the default condition, we chose 5μL (29.07 J) for the high-quality type and 2.5μL (14.535 J) for the low-quality type. This reward could represent an individual flower that offers a large reward or a small patch of several flowers, each offering smaller quantities, or it could represent a larger patch of flowers that is shared by several bees.

We tested different refill times (*t*_refill_) for food sources: 0, 1200 and 3600 ticks (Stout & Goulson, 2002). When *t*_refill_= 0, food sources became rewarding again immediately after the visit of a bee. This simulates conditions under which bees have a high probability of finding a reward after landing on food source, which might occasionally occur at food patches. With *t*_refill_= 3600, a food source (flower or patch) was empty for the equivalent of an hour after it had been visited by a bee, leading to intense exploitation competition among bees. The number of food sources per type in the simulated environment varied between 1500 (low abundance) and 4500 (high abundance). Default conditions simulated even numbers of food sources for both food source types, but we also explored uneven food source abundances (Table 1). We measured the average foraging distance of bees during a simulation run to confirm that the simulated conditions led to naturally realistic average foraging distances for many social bees (271 ± 130 m; range 63-581 m; N = 1800 simulations in default conditions) (Kohl et al., 2020; Van Nieuwstadt & Iraheta, 1996; Walther-Hellwig & Frankl, 2000).

The energy collected by agents with a full crop was estimated in the following way: *Apis mellifera* can carry up to ∼70 μL of nectar in their crop, but they usually carry less (I’Anson Price et al., 2019; Núñez, 1966). The crop load has been shown to depend on the quality of the visited food source, with lower quality food sources leading to smaller crop loads (Núñez, 1966, 1970). Agents visiting the low-quality flower type foraged until their crop contained 25μL, whereas agents visiting the high-quality food type collected 50μL per foraging trip. Generalist bees that choose indiscriminately have an intermediate crop load, reflecting the relative number of high- and low-quality food sources in the environment. For example, in an environment with an even number of high- and low-quality food sources, they collect 37.5μL per foraging trip. Sugar concentration of collected nectar varies considerably from c. 10-70% (I’Anson Price, 2018; Seeley, 1986; Willmer, 2011). We chose an average sugar concentration of 35%, providing 5.814 J/μL.

Each simulation lasted 36,000 ticks (i.e. 10 hours), simulating a day with good foraging conditions. We measured the total energy collected by a colony during this period divided by the number of agents (Energy/bee). Our main questions were if the energy/bee depended on flower constancy (vs. indiscriminate choice), communication (vs. no communication), refill time, the number of food sources and reward size. We also tested situations when flower constancy was lower after visiting a low-quality food source (*Constancy*_LQ_) (Grüter et al., 2011), when there were 4 food source types and when indiscriminate flower choice increased the time to extract a reward from a food source (*i*.*e*. to simulate cognitive constraints) (Chittka et al., 1999).

### Sensitivity analysis and model exploration

We varied a range of other factors to explore how they affected our results. These included colony size, crop load size, flower stay time, metabolic costs, nest stay time, Lévy flight *μ*, selectivity of communication (*i*.*e*. bees foraging on low-quality food source become *influencers* with the same probability as those foraging on the high-quality type) and the shape of the recruitment curve (see Fig. S1).

We performed 30 runs per parameter combination. We do not provide *p*-values due to the arbitrariness of the simulation number but indicate 95%-confidence intervals to facilitate interpretation of effect sizes.

## Results

### Food source abundance and refill speed

We found that communication about the high-quality flower type did not affect the collected energy if bees chose food sources indiscriminately (Fig. 2) (the raw data from which the figures were built can be found in the supplementary material). However, if colonies were flower constant, communication increased the energy collected by bees in all situations when the two flower types were equally abundant (Fig. 2, see also Figs. 4-8), showing an interaction between flower constancy and communication. The combination of communication and flower constancy was relatively more beneficial when high-quality food sources were easy to find, either because they were highly abundant (Fig. 2c,f,i) or because visited food sources replenished quickly (Fig. 2a,b,c).

**Fig. 2.**
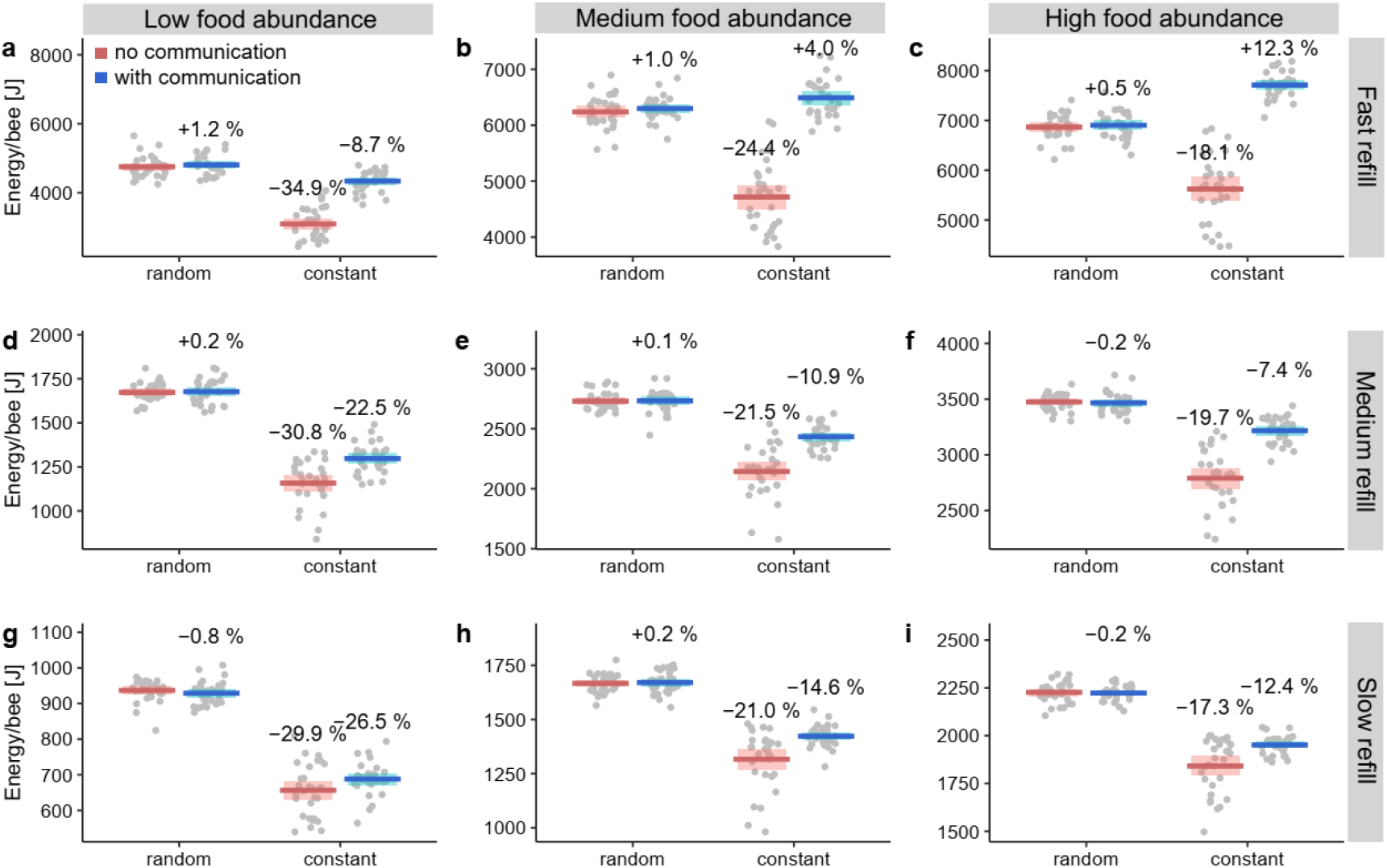
Energy collected per bee (Joule) under varying food abundances (1500, 3000 and 4500 food sources per type) and refill times (0, 1200 and 3600 ticks). Colonies either showed flower constancy (constant) or they chose food source indiscriminately (random). Plots show the mean and the 95%-confidence based on 30 simulations (grey dots). Numbers show % of change compared to random choice without communication.

In the most favourable conditions, flower constancy in combination with communication was the most successful combination (Fig. 2b, c). In all other conditions, indiscriminate choice was the most successful strategy.

The relative abundance of the two food source types also played an important role. Flower constancy combined with communication was relatively more successful when high-quality food sources were more common than the low-quality flower type compared to when they were rarer than the lower-quality flower type (Fig. 3). When high-quality food sources represented the common flower type, colonies with flower constancy and communication were either more successful (Fig. 3b) or not much less successful than colonies with indiscriminate choice (Fig. 3d).

**Fig. 3.**
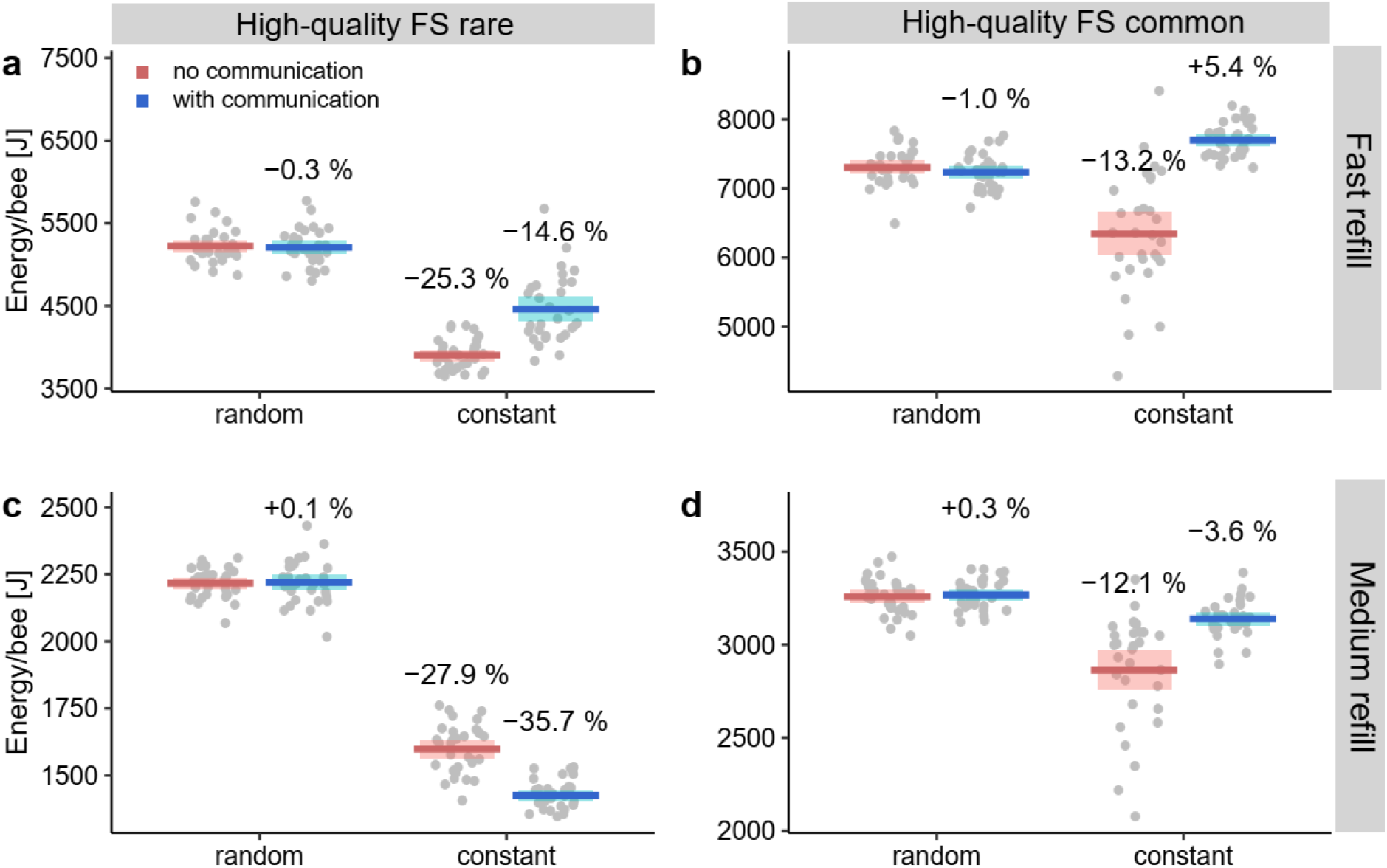
Energy collected per bee (Joule) when high-quality food sources were either rare (1500) or common (4500) compared to the low-quality food sources (3000). Default values were used for the other parameters (Table 1).

However, indiscriminate choice was considerably more successful when high-quality food sources were in the minority (Fig. 3a,c). When high-quality food sources were particularly difficult to find, communication lowered the foraging success of flower constant colonies (Fig. 3c). Under these circumstances, communication directs the foragers of a colony towards a rare food source, leading to long search times.

### Reward sizes

Reward quantities are known to affect flower constancy, with bees becoming more flower constant with increasing reward quantities (Chittka et al., 1997; Grüter et al., 2011; Wells & Rathore, 1994). In accordance with this observation, we found that flower constancy became relatively more successful (energy/bee) as reward sizes of both high- and low-quality food sources increased (Fig. 4). However, indiscriminate choice was the most successful strategy in many tested environments.

**Fig. 4.**
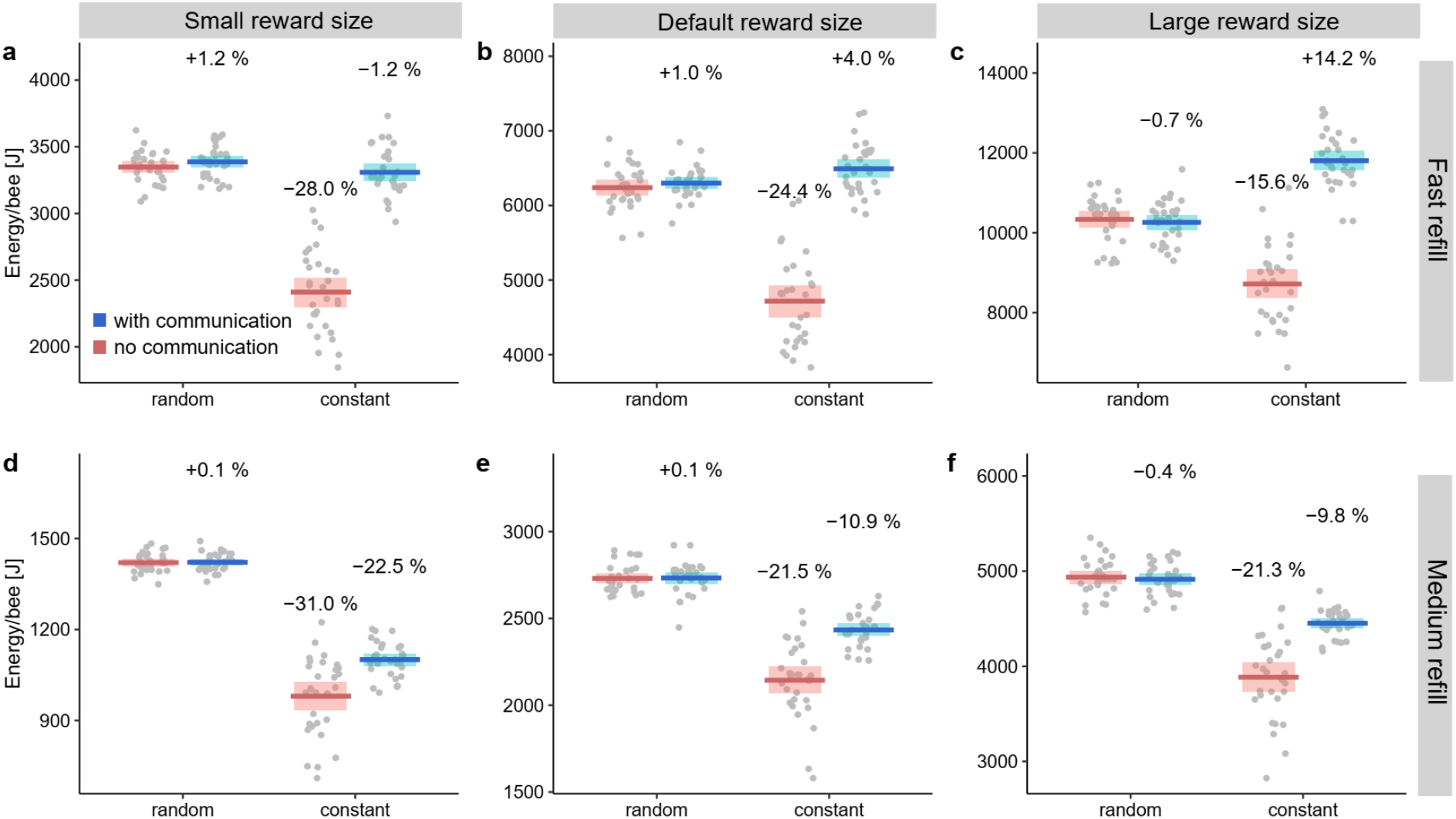
Energy collected per bee (Joule) when rewards were (a) smaller (2.5μl and 1.25μl) or (c) larger (10μl and 5μl) than in the (b) default situation (5μl and 2.5μl). Medium food source abundance was simulated; default values were used for the other parameters (Table 1). Blue shows means and 95%-CI for colonies using communication, red shows data for colonies choosing indiscriminately.

### Time needed to collect a reward

The time bees need to extract a reward from a flower will affect the time costs of foraging decisions and, if the refill time is >0, it will affect the number of depleted food sources in the environment. Under default conditions, bees needed 60 ticks (1 minute) to obtain the reward from a flower/food patch. We explored how different values for *t*_Flower-stay_ affected the benefits of flower constancy and communication. Increasing the time needed to obtain a reward increased the relative benefits of combined flower constancy and communication compared to short reward collection times (Fig. 5).

**Fig. 5.**
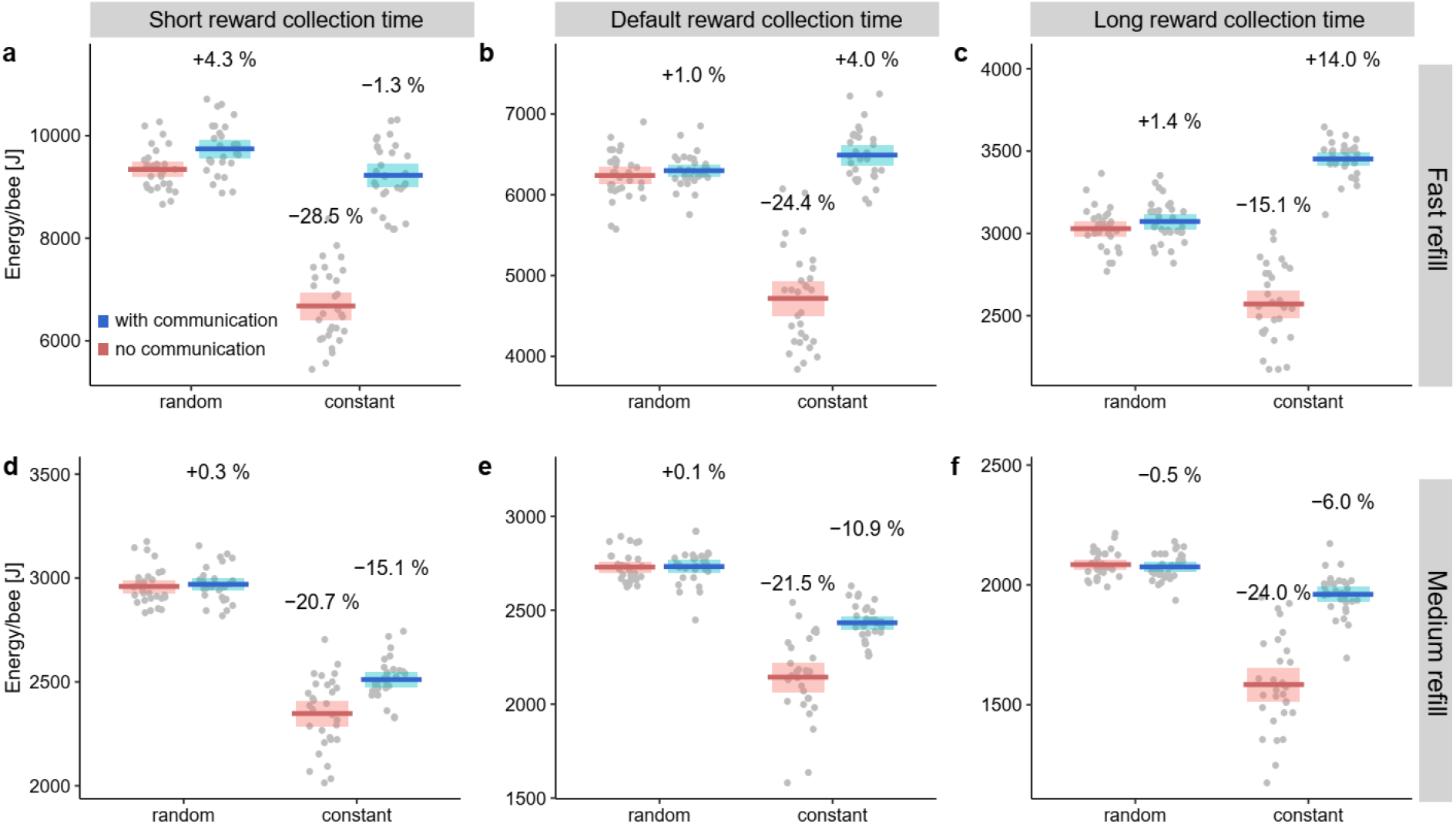
Energy collected per bee (Joule) when the time required to obtain a reward from a food source was (a,d) shorter (20 ticks) or (c,f) longer (180 ticks) than in the (b,e) default situation (60 ticks). Default values and a medium food source abundance were simulated (Table 1).

### Quality dependent flower constancy

Under default conditions, flower constancy did not depend on the quality of the food source (“spontaneous flower constancy”, Hill et al., 1997). We simulated situations when bees visiting a low-quality food source were slightly less flower constant (they had a 90% or a 95% chance to remain flower constant on the subsequent visit, as in Grüter et al., 2011). Our results show that this quality-dependent flower constancy considerably improves the energy collected by colonies following this strategy of quality-dependent flower constancy (Fig. 6).

**Fig. 6.**
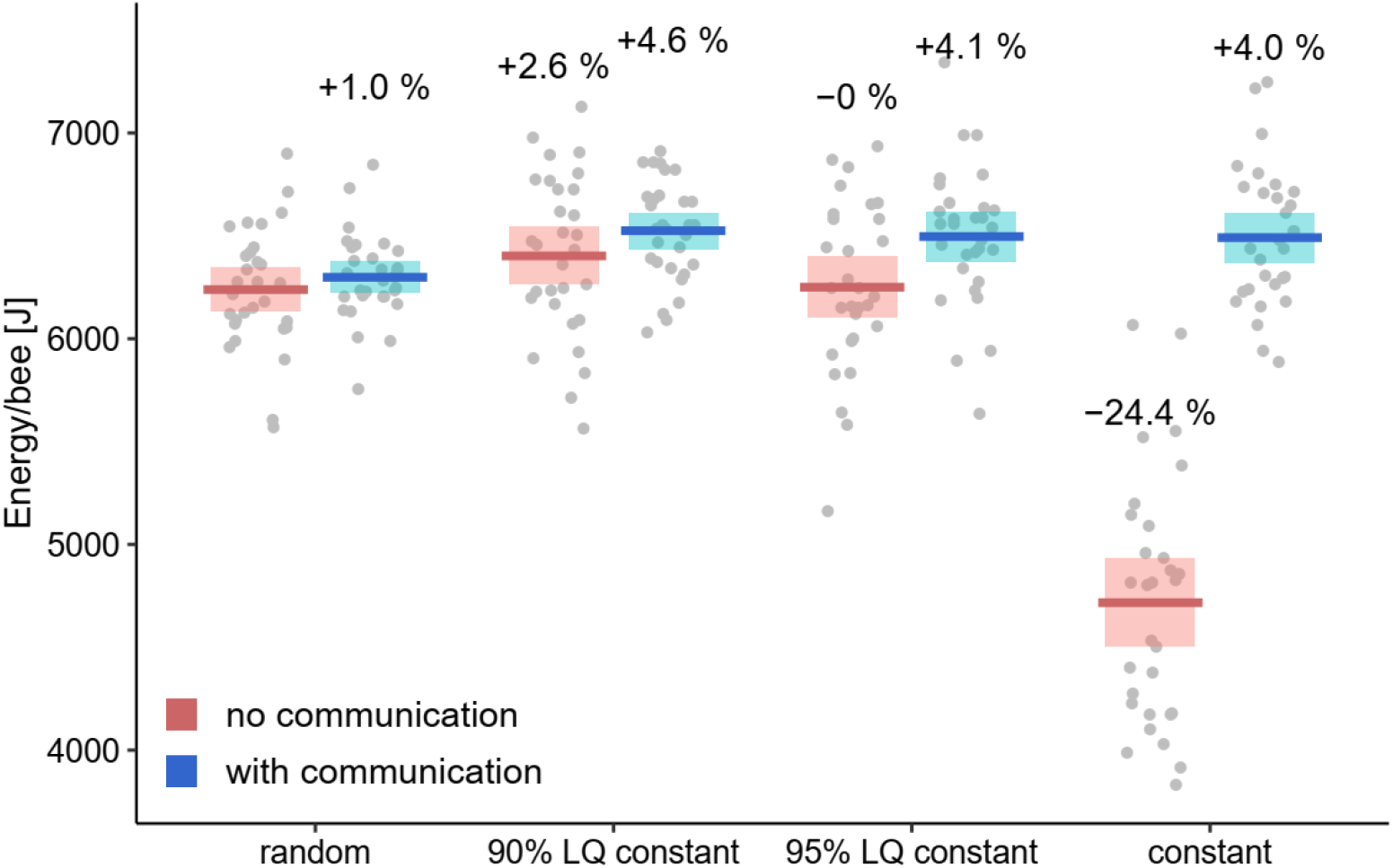
Energy collected per bee (Joule) when bees foraged indiscriminately, when they showed reduced flower constancy after visiting a low-quality food source (90% LQ constant or 95% LQ constant) and when they were strictly flower constant. Refill time was 0, default values were used for the other parameters (Table 1).

### Exploring environments with 4 food types

When environments provide four different types of food sources rather than two, flower constancy is less favourable overall (Fig. 7). In other words, indiscriminate flower choice is highly beneficial in an environment where flower constancy would limit the options a forager has to a small subset (25% of all food sources) of all available food sources than with two food source types (Fig. 7).

**Fig. 7.**
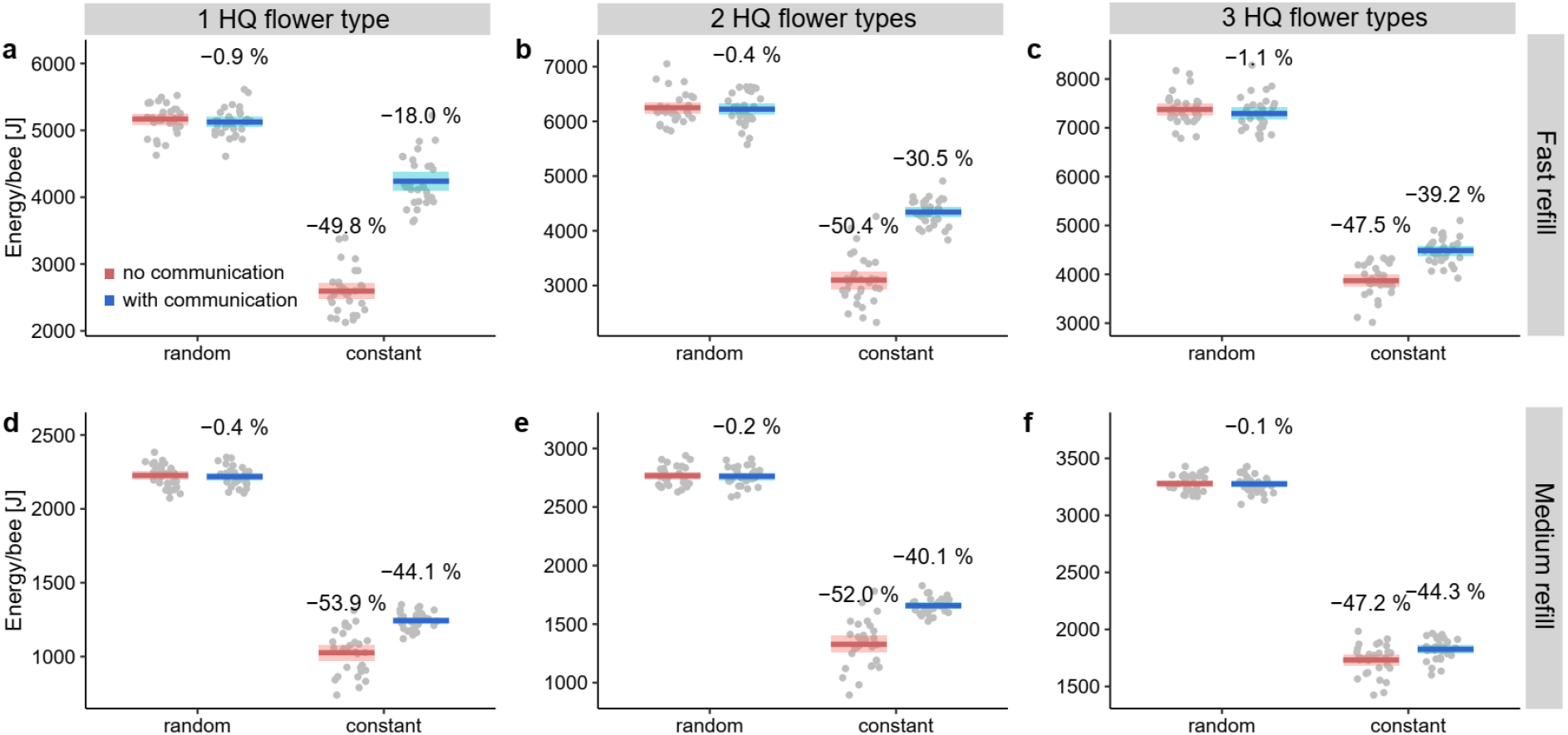
Energy collected per bee (Joule) when bees foraged in an environment of four flower species. In (a)& (d), one of the four plant types was of high quality, while the remaining three types were of low quality. In (b) & (e), two of four types were of high quality and in (c) & (f), three of flower types were of high quality. Food sources were refilling either at a fast or a medium rate.

We tested situations where one, two or three of the four plant types were of high quality, while the remaining food sources were of low quality. While it was always beneficial to use communication when colonies were also flower constant, the relative benefits of communication diminished as the number of high-quality food types and the refilling time increased. Unsurprisingly, therefore, foraging in an environment that consists mainly of high-quality food sources belonging to different plant species somewhat diminishes the value of using communication to direct foragers towards higher-quality food sources.

### Time penalty for non-specialists

So far, we have assumed that there are no additional time costs (e.g. as a result of cognitive limitations) for bees that do not specialise on a particular type of food source. To explore the consequences of cognitive limitations, we simulated situations when indiscriminate bees require more time to extract a reward from a food source compared to flower constant bees. A time penalty for indiscriminate bees favours flower constant colonies, especially those that also communicate the high-quality flower type to nestmates (Fig. 8).

**Fig. 8.**
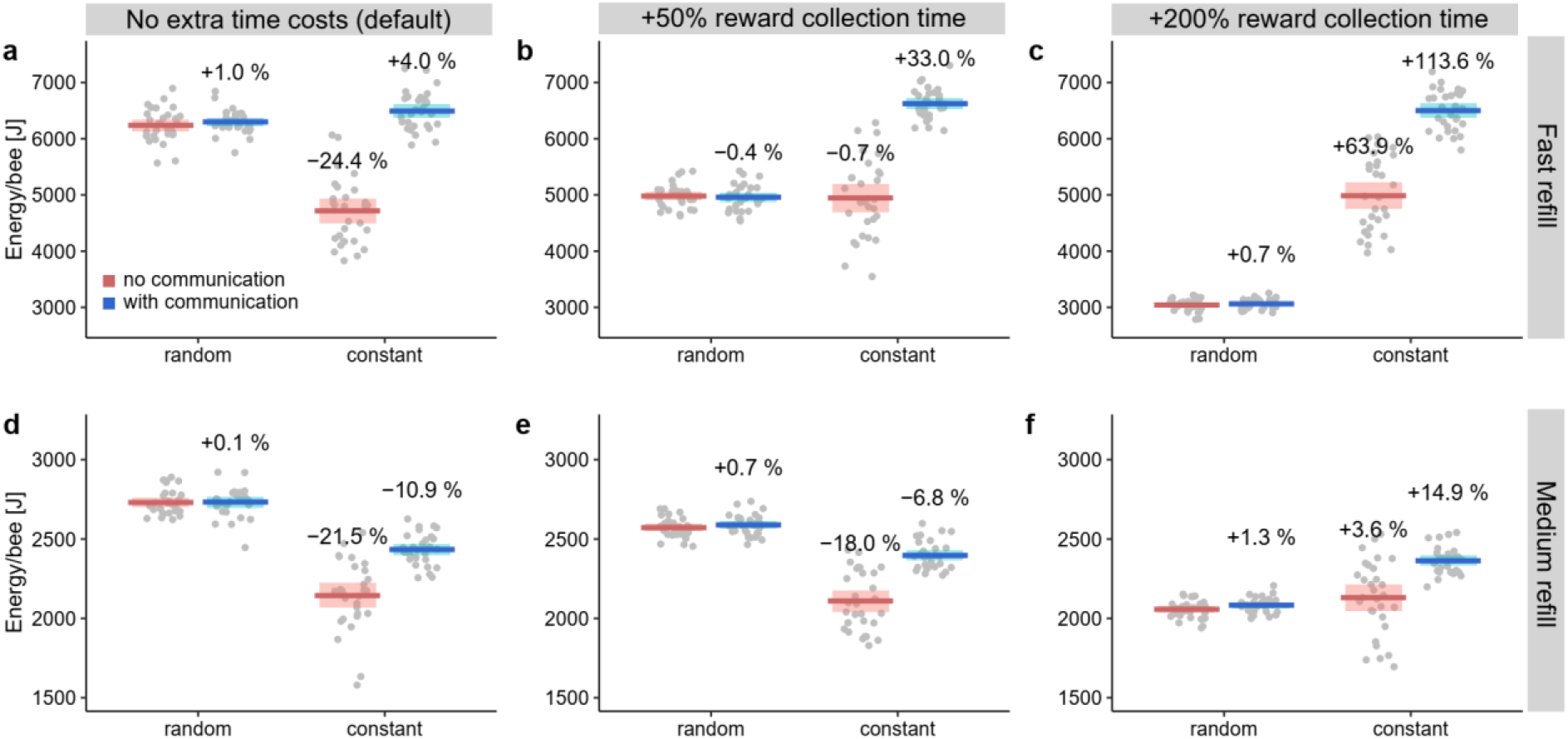
Energy collected per bee (Joule) when non-specialists needed 50% (b, e) or 200% (c, f) more time than flower constant bees to obtain a reward from a food source. Food sources were refilling either at a fast (*t*_refill_ =0) or a medium rate (*t*_refill_ = 1200). Default values were used for the other parameters (Table 1).

### Sensitivity analysis and model exploration

Varying colony size from 5 to 300 (Fig. S2) does not greatly affect the general pattern observed for the default colony size of 100 (see Fig. 2). When food sources refill at a fast rate (Fig. S2a), bees do not experience exploitation competition and colony size does not affect the energy collected by individual bees. Increasing the refill time while also increasing the number of agents searching for food, on the other hand, increases exploitation competition and, therefore, lowers the energy collected by individual bees (Fig. S2b,c).

Using different recruitment curves (Fig. S1) had no noticeable effect on the energy collected by bees, but non-selective recruitment (recruitment to both high- and low-quality food sources) lowers the collected energy to levels similar to those of flower constant colonies without communication (Fig. S3). Changing the metabolic costs of flying or the Lévy-flight *μ* has little effect on the overall pattern (Fig. S4, S5), whereas increasing the time spent inside the nest in-between foraging trips reduces the energy collected by bees, but less so in colonies with flower constancy (Fig. S6). Thus, longer nest stay times favour flower constancy. Flower constancy was also favoured when bees had smaller crop sizes (Fig. S7).

## Discussion

Results from our simulations suggest that flower constancy without communication is less successful than indiscriminate choice under all tested conditions. Flower constancy imposes significant costs because it (*i*) limits the available options to a subset of all available flowers, thereby increasing time and energy costs during foraging, and (*ii*) causes many foragers to specialise on a sub-optimal flower type. Communication about the high-quality flower type positively interacted with flower constancy (Fig. 2) and considerably improved the foraging success of flower constant colonies by. Communication allows a colony to focus on high-quality flowers, thereby reducing the second type of cost (*ii*). Many species of social bees have evolved mechanisms of reward-quality dependent recruitment communication, which allow *influencers* to affect the foraging decisions of their nestmates towards a particular flower type, mainly via olfactory learning (Dornhaus & Chittka, 1999; Farina et al., 2012; Jarau & Hrncir, 2009; Lindauer & Kerr, 1960; von Frisch, 1967). This, in turn, lowers the benefits of sampling alternative flower species and highlights the importance of social information use as a process of information-filtering (Grüter et al., 2010; Rendell et al., 2010). Our findings can help explain why social bees tend to be more flower constant than solitary bees (e.g. Smith et al., 2019; Waser, 1986).

The general foraging conditions had a strong effect on the value of flower constancy and the strength of its interaction with communication. Flower constancy in combination with communication was the most successful strategy when foraging conditions were very favourable, while indiscriminate choice was the better strategy when foraging options were more limited. For instance, flower constancy in combination with communication was beneficial when foragers did not encounter empty food sources (refill time of 0) and food sources were abundant (Fig. 2b,c), when most food sources were of high-quality (Fig. 3b) and when rewards were large (Fig. 4c). These findings are consistent with empirical studies showing that bees are more flower constant when flower density is higher (Chittka et al., 1997; Kunin, 1993; Marden & Waddington, 1981) and rewards are larger (Chittka et al., 1997; Greggers & Menzel, 1993; Grüter et al., 2011). Similarly, predator-prey models show that the abundance of a prey item has a positive effect on diet specialisation of the predator (Pulliam, 1974). If food sources took time to replenish, resulting in many empty food sources due to exploitation competition, indiscriminate choice was more successful (Fig. 2d-i), suggesting that rejecting flowers due to flower constancy is more costly in environments that offer fewer options.

Changes in the temporal dynamics of foraging trips affected the performance of the different strategy by changing the relative costs of ignoring flowers (*i*) and choosing suboptimal food sources (*ii*). Flower constancy in combination with communication performed relatively better if bees required more time to extract a reward from a food source (Fig. 5). One explanation for this is that visiting low-quality food sources, which is common with indiscriminate choice, becomes relatively more costly as the time costs of a flower visit increase. Thus, longer flower handling times, e.g. due to a complex flower morphology, favour flower constancy from both an adaptive and a constraints-based perspective (see Chittka et al., 1999 for arguments based on cognitive constraints). Flower constancy in combination with communication performed relatively better when bees stayed in their nest longer (Fig. S6) and had smaller crops (Fig. S7). These findings are somewhat puzzling, but one explanation could be that longer nest stay times provide *influencers* with more opportunities to communicate their findings to other bees. When food sources need time to replenish, longer nest stay times will reduce the number of depleted food sources a bee encounters, which favours flower constancy in combination with communication (Fig. 2b,c). Similarly, when bees have smaller crop loads, they visit fewer food sources per trip and spend a larger proportion of their time in the nest, reducing exploitation competition and the number of depleted food sources. Crop size will depend on body size and one might, therefore, predict that smaller bees are more flower constant, which is consistent with comparative data (Smith et al. 2019). However, it is unlikely that there is straight forward relationship between crop size, body size and flower constancy in nature because body size covaries with numerous other extrinsic and intrinsic factors, including foraging conditions, metabolic costs, flying speed or sensory acuity (Gervais et al., 2020; Grab et al., 2019; Spaethe et al., 2007), all of which might affect flower constancy.

The Western honey bee *Apis mellifera* is strongly flower constant, but there is disagreement about whether and when flower constancy depends on the profitability of visited flowers. Some studies have suggested that flower constancy is often “spontaneous”, *i*.*e*. unrelated to reward size (Wells & Wells 1983; Hill et al. 1996, 2001; Sanderson et al. 2006), whereas others have found that honey bees adjust flower constancy according to the profitability of rewards (Greggers & Menzel 1993; Chittka et al. 1997; bumble bees: Heinrich 1976, 1979b; reviewed in Grüter & Ratnieks 2011). Our simulations show that context-dependent flower constancy is more successful than strict (“spontaneous”) flower constancy (Fig. 6). When bees visiting the less profitable food type were only 90-95% flower constant, colonies collected about 25% more energy than colonies with strict flower constancy. As is the case with communication, context-dependent flower constancy allows bees to switch from the low-quality to the high-quality flower species over time (type (*ii*) costs).

Human impacts have significantly affected the diversity of plant species found in some environments, especially in intensively farmed habitats (e.g. Potts et al., 2010; Tew et al., 2021), which is likely to affect the costs and benefits of flower constancy. In our simulations, flower constancy performed considerably worse when there were four rather than two flower types (Fig. 7). With more plant species present, flower constant bees will ignore most of the available options and focus on a small subset of all food sources, thereby dramatically increasing opportunity costs (type (*i*) costs). Thus, bees should be less flower constant in more diverse foraging environments. Flower constant bees, in turn, might suffer a reduction in foraging success in more biodiverse habitats. These findings challenge the reasoning behind the “costly information hypothesis”, which argues that flower constancy is an adaptive foraging strategy because acquiring information about suitable alternatives would cost too much time and energy if there are several plant species available (Chittka et al., 1999; Grüter & Ratnieks, 2011). In flower diverse environments, bees should accept even low-quality food source if it means they can cut time and energy costs imposed by flower constancy. Empirical studies on the links between floral diversity and flower constancy provide contrasting results. While Gervais et al. (2020) and Martínez-Bauer et al. (2021) found that increasing plant diversity was associated with lower flower constancy in *Bombus impatiens* and *B. terrestris*, Austin et al. (2019) found that bumble bees became more flower constant when there are more options available. The latter finding is more consistent with a “cognitive limitations” perspective, since deciding among more options would be cognitively more challenging and flower constancy, therefore, a possible solution to avoid switching costs (see also Chittka et al. 1997; Gegear & Thomson 2004). Decision making is often impaired as the number of choices increases (Latty & Trueblood, 2020). The different studies differ in that the first two were performed under natural conditions, whereas Austin et al. (2019) was experimental. Non-experimental surveys can be confounded by numerous factors, such as differences in rewards, clustering of flowers or management, whereas experimental studies might fail to capture crucial features of natural environments that affect decision-making (Fawcett et al., 2014).

Agent-based models have important limitations. Simulation outcomes depend on the underlying assumptions and the parameters chosen when building the model, some of which are arbitrary or simplistic. As a result, ABMs potentially miss important natural features that shape decision-making (Fawcett et al., 2014). For example, we assumed that food sources are randomly distributed, whereas natural foraging environments are often spatially heterogenous and patchy, which is likely to affect the value of flower constancy. Patchiness can lead to flower constancy “by accident” if bees forage in large patches, even if they choose flowers indiscriminately. We might, therefore, expect increasing patchiness to lead to more similar outcomes for flower constant and indiscriminate foragers. Pulliam’s (1974) predator-prey model found that an increasingly clumped prey distribution favours a more specialised diet in predators and we might expect a similar finding in plant pollinator interactions. Agent-based models also have important strengths because they allow us to systematically vary factors that cannot be manipulated experimentally, such as tuning flower constancy or recruitment communication while keeping all other factors constant. ABMs should be seen as a useful tool to complement empirical studies.

One of the aims of our model was to test whether flower constancy could be an adaptive strategy *per se* under some foraging conditions, *i*.*e*. in the absence of cognitive constraints. If, however, switching between flower species leads to increased time costs or reduced reward sizes (Chittka et al., 1999; Darwin, 1876; Grüter & Ratnieks, 2011; Lewis, 1986; Raine & Chittka, 2007), flower constancy, with or without communication, becomes a much more beneficial strategy under a wide range of conditions (Fig. 8c,f). The reasons for flower constancy in pollinators are likely to be complex and depend on both constraints and adaptive processes, to varying degrees in different species. However, our results suggest that a more pronounced flower constancy in social bees is more likely due to increased performance of flower constancy in social species due to social traits, rather than the result of poorer cognitive abilities in social bees compared to solitary bees (Amaya-Márquez & Wells, 2008; Dukas & Real, 1991).

## Supporting information

Raw data with 2 flower types

Raw data with 4 flower types

Main model code for NetLogo

## Acknowledgements

We thank the University of Bristol Ecology group for comments and feedback on the results of this study.

---

**Fig. S1.**
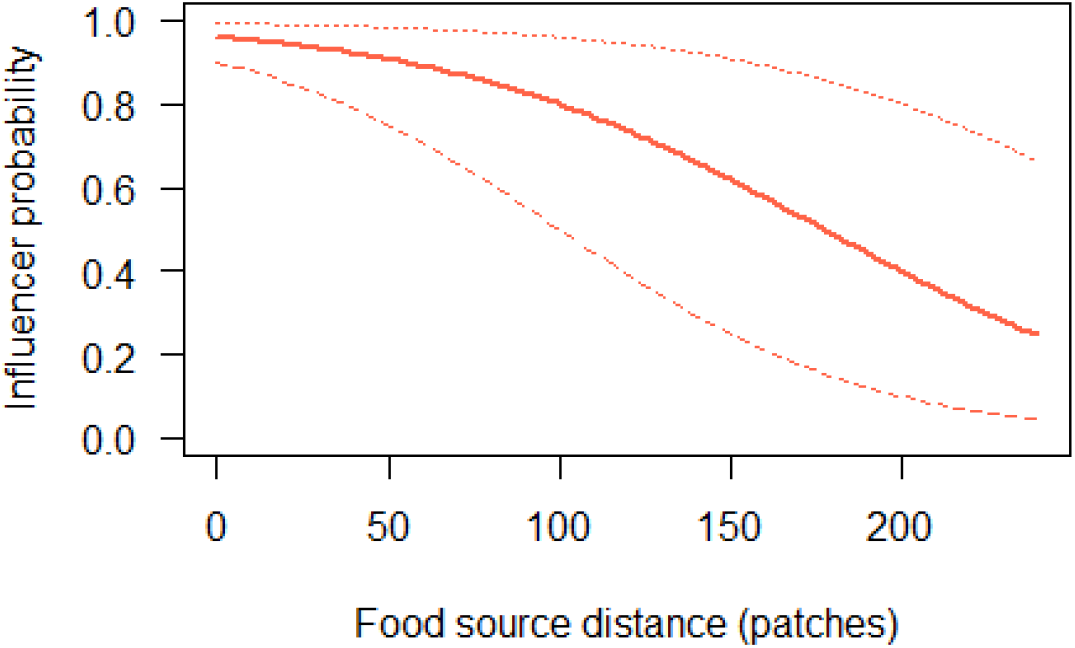
Probability that a returning bee that visited the high-quality food type (red) becomes an influencer inside the nest. Bees visiting the low-quality food type did not become influencers under default conditions. Thick red line shows the default probability. The other two lines show other tested probability curves. 1 patch ∼ 5 m.

**Fig. S2.**
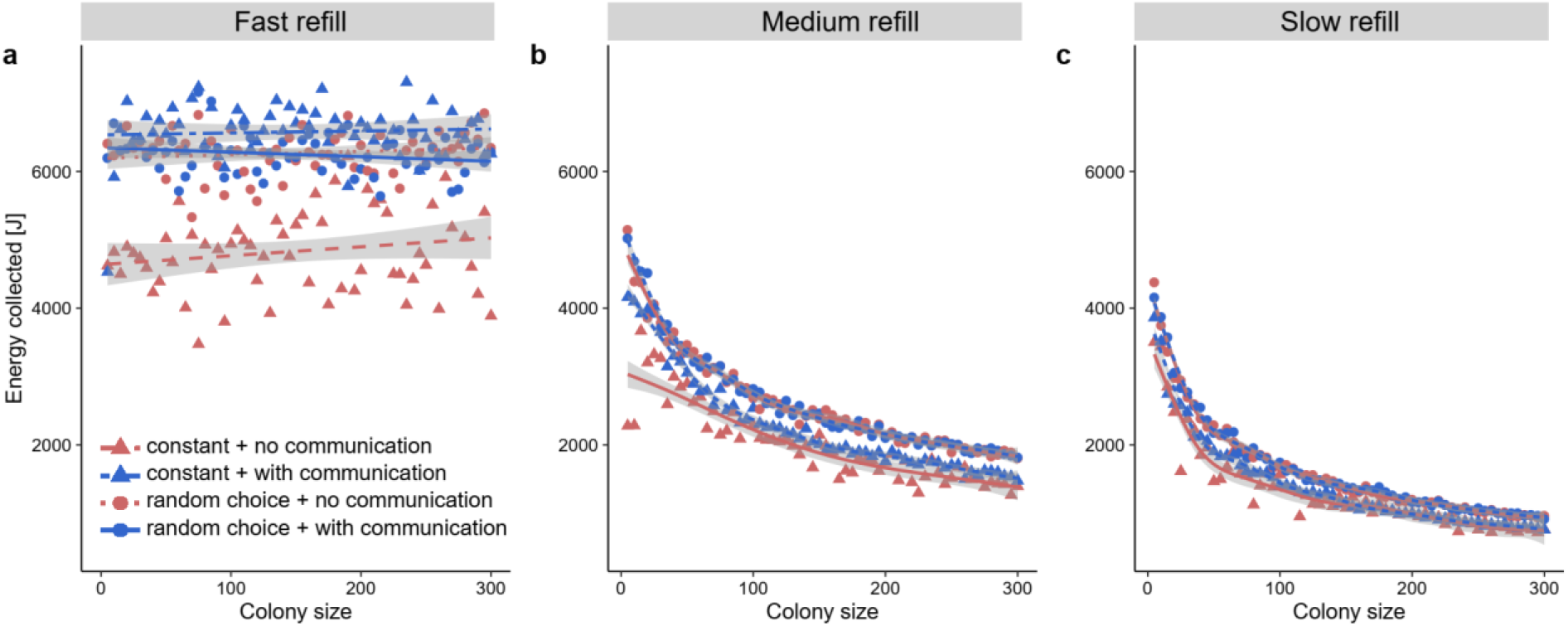
Energy collected per bee for different colony sizes and different refill speeds (fast refill, *t*_refill_ = 0; medium refill, *t*_refill_ = 1200; slow refill, *t*_refill_ = 3600). Lines in (a) show best fit lines based on linear regression. In (b) and (c), generalised additive models (GAM) were used to fit curves. Grey bands show 95% confidence intervals. Default values were used for other parameters.

**Fig. S3.**
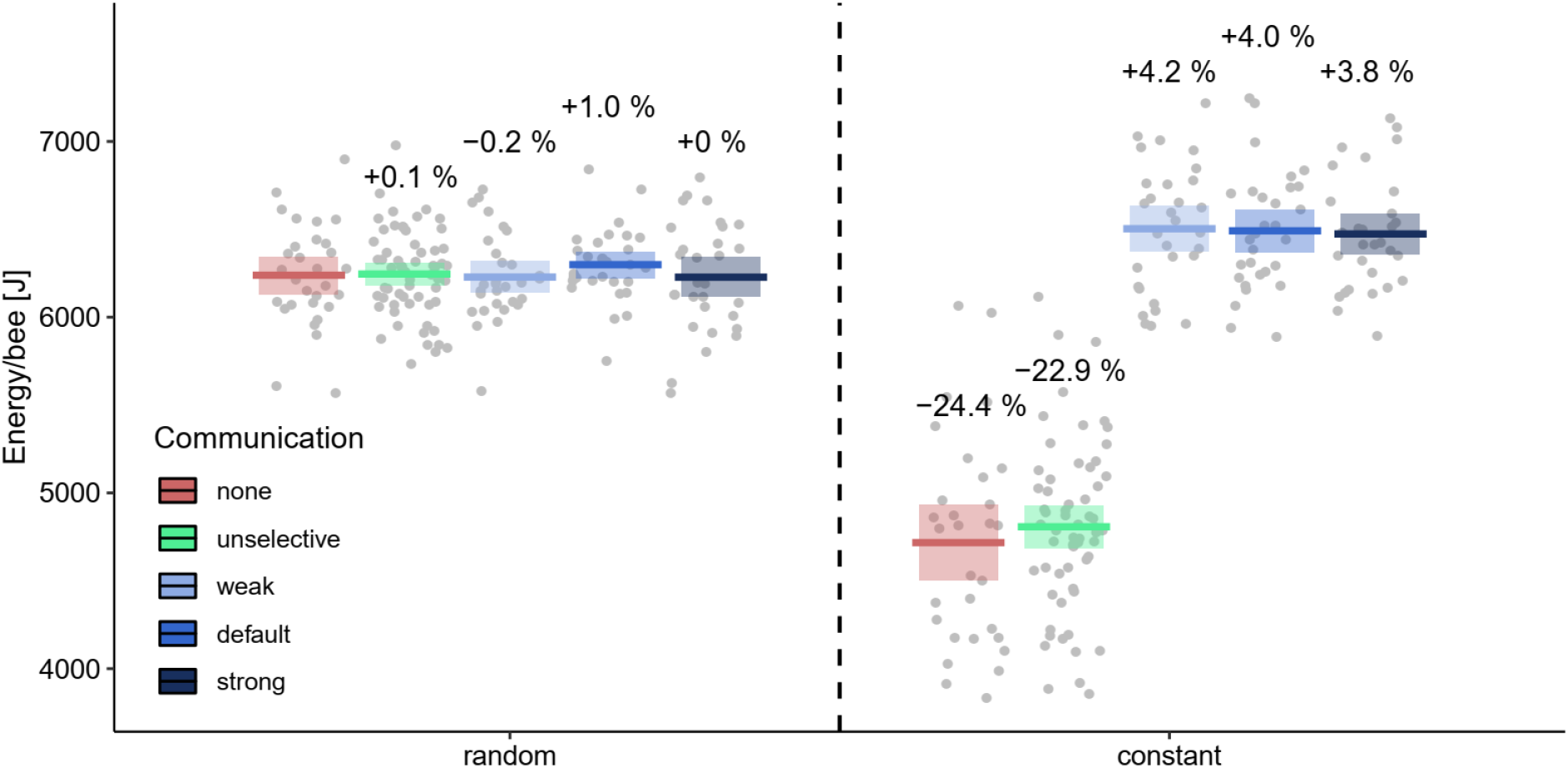
Energy collected per bee for different types of communication, with or without flower constancy. The different recruitment strengths (weak, default and strong) correspond to the three different curves shown in S1. Unselective refers to a situation where bees communicate equally about high- and low-quality food sources (assuming the default curve shown in Fig. S1). Means and 95%-confidence intervals are shown.

**Fig. S4.**
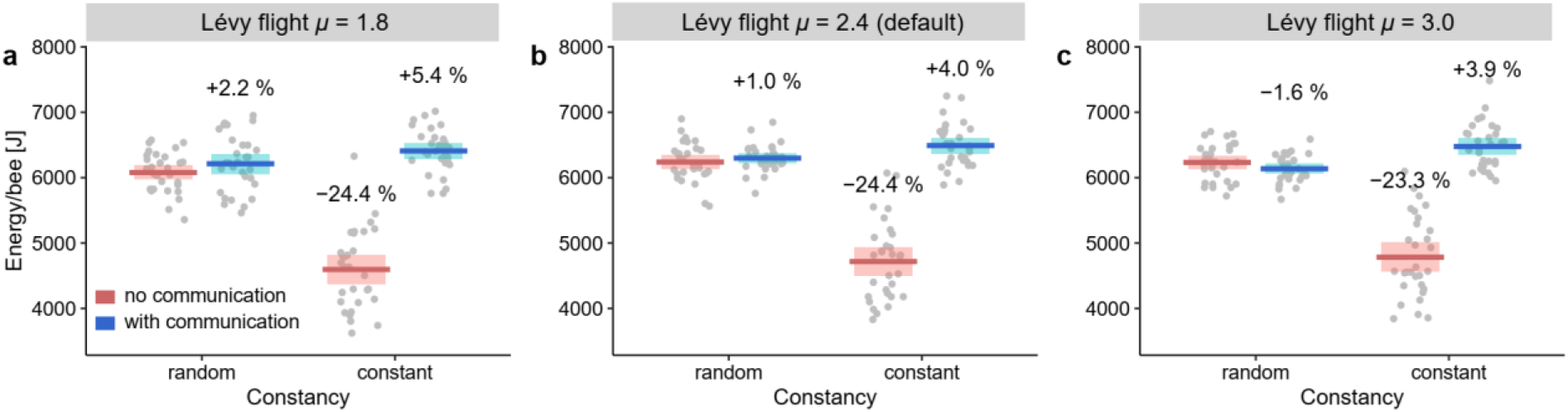
Energy per bee (Joule) collected using different Lévy-flight *μ* values, which determines the distribution of the segment lengths *l* that constitute a Lévy-flight, with *P*(*l*)∼*l*^-μ^. Food sources that refill immediately were simulated. Default values were used for the other parameters (Table 1).

**Fig. S5.**
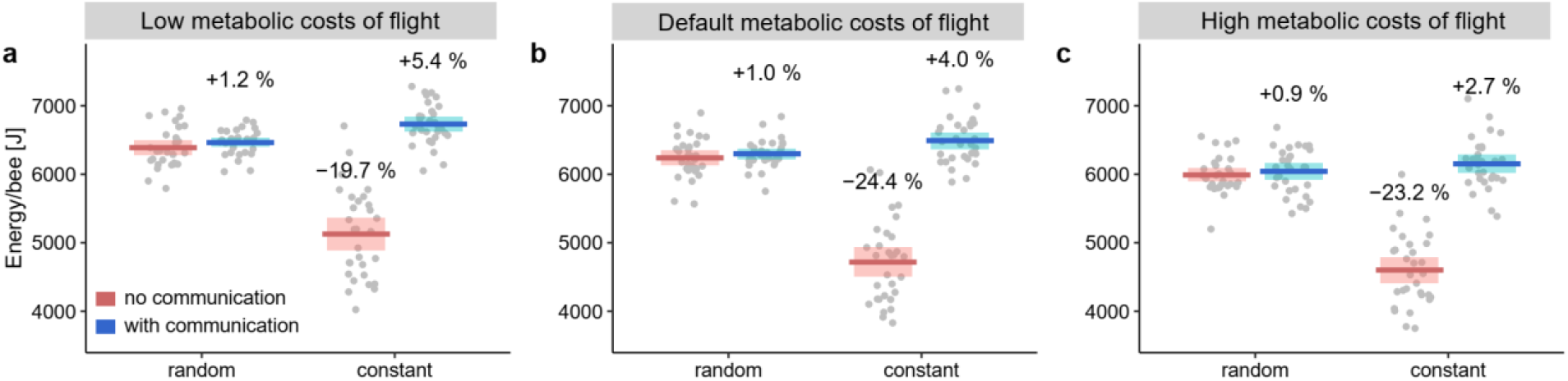
Energy per bee (Joule) collected using different flight energy costs (low = 0.016 J/tick; default = 0.032 J/tick; high = 0.064 J/tick). Food sources refilled immediately. Default values were used for the other parameters (Table 1).

**Fig. S6.**
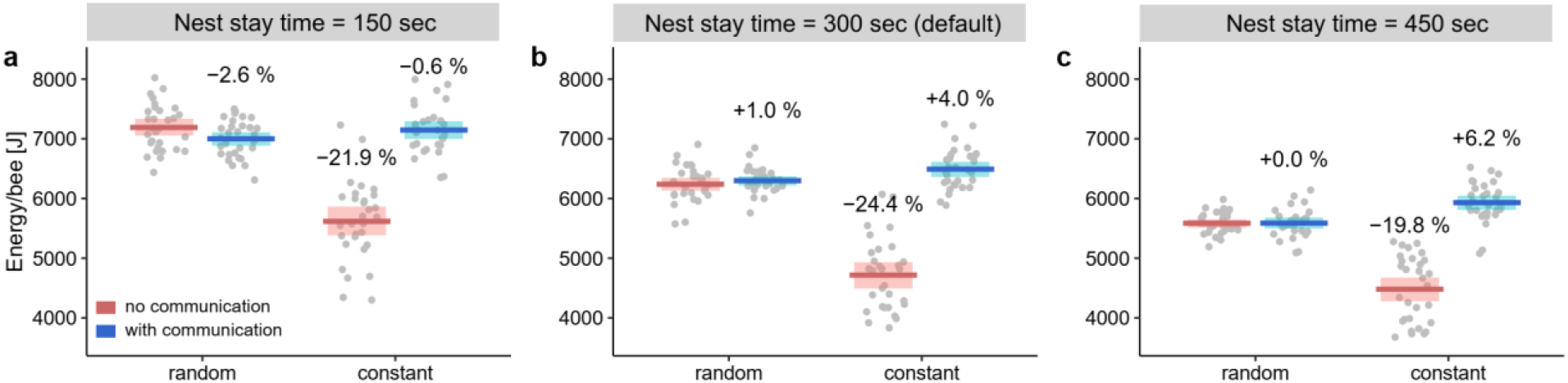
Energy per bee (Joule) collected using different nest stay times. Food sources refilled immediately. Default values were used for the other parameters (Table 1).

**Fig. S7.**
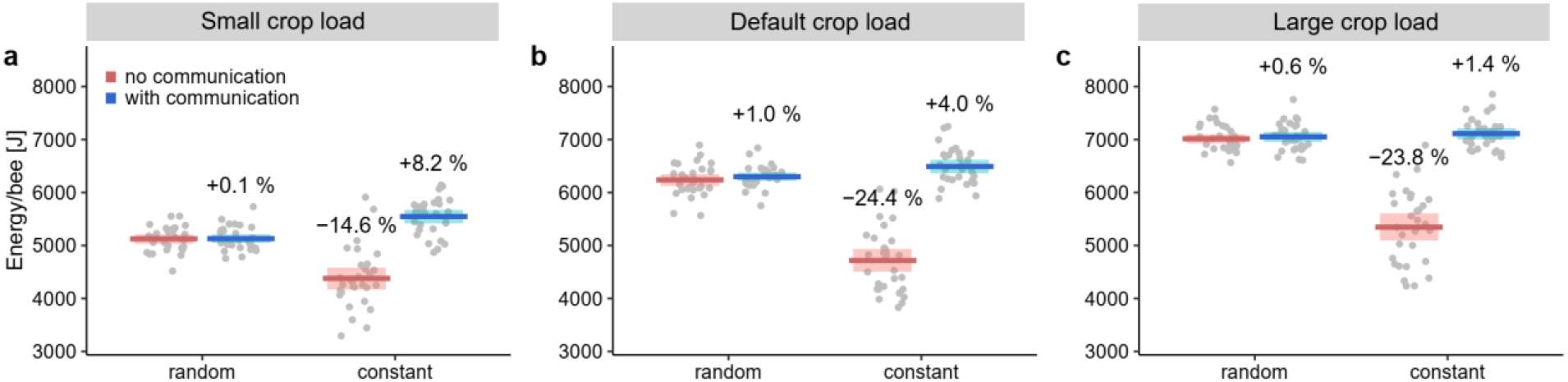
Energy per bee (Joule) collected with different crop loads. (a) Small crop loads mean that bees constant to high-quality food sources filled their crop to 25μl, whereas bees constant to low-quality food sources returned to their nest if their crop load was 12.5μl. Bees choosing food sources randomly had intermediate crop loads (18.75μl). (c) Bees with large crop loads foraged until they had 100μl (constant to high-quality food sources), 50μl (constant to low-quality food sources or 75μl (randomly choosing bees). Food sources refilled immediately. Default values were used for the other parameters (Table 1).

