## Supplementary material for "When should bees be flower constant? An agent-based model highlights the importance of social information and foraging conditions": Main model code for NetLogo

**Interface**

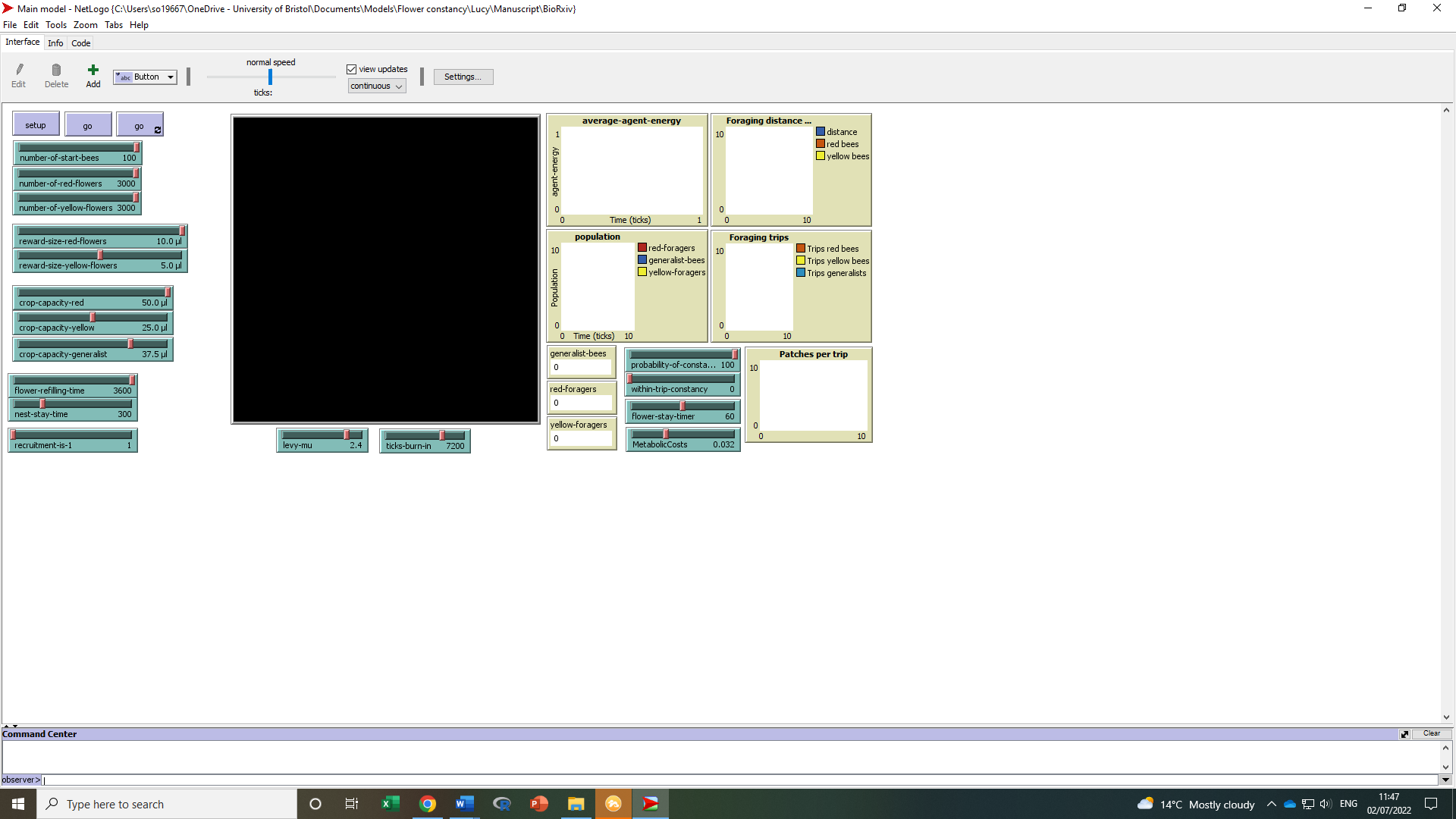

**Model code**

globals

[ nest-energy

foraging-trips-red-bees

foraging-trips-yellow-bees

foraging-trips-generalist-bees

foraging-distance	;; used for foraging distance

foraging-distance-red	;; used for foraging distance

foraging-distance-yellow	;; used for foraging distance

total-visits	;; used for foraging distance

total-visits-red	;; used for foraging distance

total-visits-yellow	;; used for foraging distance

logisticA	;; for logistic communication function, see Railsback p. 215-216.

logisticB	;; for logistic communication function

total-patches	;; used for patches per trip

]

breed [generalist-bees generalist-bee]

breed [red-flowers red-flower]

breed [yellow-flowers yellow-flower]

breed [red-foragers red-forager]

breed [yellow-foragers yellow-forager]

breed [return-red-foragers return-red-forager]

breed [return-yellow-foragers return-yellow-forager]

breed [return-generalist-foragers return-generalist-forager]

breed [inside-nest-workers-red inside-nest-worker-red]

breed [inside-nest-workers-yellow inside-nest-worker-yellow]

breed [inside-nest-workers-generalist inside-nest-worker-generalist]

breed [empty-red-flowers empty-red-flower]

breed [empty-yellow-flowers empty-yellow-flower]

breed [red-influencers red-influencer]

breed [yellow-influencers yellow-influencer]

breed [red-stop-foragers red-stop-forager]

breed [yellow-stop-foragers yellow-stop-forager]

breed [return-stop-foragers-red return-stop-forager-red]

breed [return-stop-foragers-yellow return-stop-forager-yellow]

breed [return-stop-foragers-generalist return-stop-forager-generalist]

turtles-own [

forager-energy

inside-nest-stay-time

empty-flower-time

flower-stay-time

trip-red

trip-yellow

trip-generalist

patch-distance	;;;;;;;;;;; for foraging distance: distance of patch

patch-distance-red

patch-distance-yellow

flower-visits	;;;;;;;;;;; for foraging distnace: total patch visits

flower-visits-red

flower-visits-yellow

flight-length	;;;;;;;;;;; for Levy flight

distance-red	;;;;;;;;;;; for communication: distance of last visited patch

distance-yellow	;;;;;;;;;;; for communication: distance of last visited patch

patches-visited-per-trip	;;; to count patches visited per trip

]

patches-own [

nest-scent

nest?

]

to setup

clear-all

ask patches [

setup-nest

setup-logcurve	;;; new for log function

]

;;;; CREATE FLOWERS ;;;;

create-red-flowers number-of-red-flowers [

setxy random-xcor random-ycor

set color red

set shape "flower"

set size 10

set flower-visits 0

set flower-visits-red 0

set empty-flower-time flower-refilling-time

set patch-distance distancexy 0 0

set patch-distance-red distancexy 0 0

]

create-yellow-flowers number-of-yellow-flowers [

setxy random-xcor random-ycor

set color yellow

set shape "flower"

set size 10

set flower-visits 0

set flower-visits-yellow 0

set empty-flower-time flower-refilling-time

set patch-distance distancexy 0 0

set patch-distance-yellow distancexy 0 0

]

;;;;;;;; CREATE BEES ;;;;

create-generalist-bees number-of-start-bees [

setxy 0 0

set size 6

set color white

set shape "butterfly"

set forager-energy 0

set flight-length 0	;;;;;;;;;; for Levy flight

set patches-visited-per-trip 0

]

reset-ticks

end

;;;;;;;;;; CREATE NEST ;;;;

to setup-nest

set nest? (distancexy 0 0) < 5

set nest-scent 300 - distancexy 0 0

set nest-energy 0

set total-patches 0

set total-visits 0

ifelse nest?

[ set pcolor white ]

[ set pcolor green ]

end

to setup-logcurve ;; setup the variables for the logistic curve ;;;;;;;;; new for log curve

let D ln (0.8 / (1 - 0.8))	; default 0.8

let C ln (0.4 / (1 - 0.4))	; 0.4

set logisticB (D - C) / (100 - 200)

set logisticA D - (logisticB * 100)

end

;;;; GO PROCEDURE ;;;;

to go

;;;;;;;;;;;;;;;;;;;;;; GO PROCEDURE ALL FLOWERS ;;;;;;;;;;;;;;;;;;;;;;

ask red-flowers [

be-eaten-red

set color red

set shape "flower"

set size 10

]

ask yellow-flowers [

be-eaten-yellow

set color yellow

set shape "flower"

set size 10

]

ask empty-red-flowers [

record-patch-distance

record-patch-distance-red

record-flower-visit

record-flower-visit-red

set color red

set shape "x"

set size 10

set empty-flower-time empty-flower-time - 1

if empty-flower-time < 0

[set breed red-flowers

set patch-distance distancexy 0 0

set patch-distance-red distancexy 0 0

reset-times ]]

ask empty-yellow-flowers [

record-patch-distance

record-patch-distance-yellow

record-flower-visit

record-flower-visit-yellow

set color yellow

set shape "x"

set size 10

set empty-flower-time empty-flower-time - 1

if empty-flower-time < 0

[set breed yellow-flowers

set patch-distance distancexy 0 0

set patch-distance-yellow distancexy 0 0

reset-times ]]

;;;;;;;;;;;;;;;;;;;;;;;;;;;;;;;;;; CREATE FORAGERS ;;;;;;;;;;;;;;;;;;;;;;;;;;;;;;;;;;

ask generalist-bees [

set size 6

set color white

set shape "butterfly"

set flower-stay-time flower-stay-timer

move-levy-flight

eat-red-flower

eat-yellow-flower

if forager-energy > (crop-capacity-generalist * 5.814) ;; 25 mu liter is 145.35 J

[set breed return-stop-foragers-generalist]

]

ask red-foragers [

move-levy-flight

eat-red-flower

set flower-stay-time flower-stay-timer

set size 6

set color red

set shape "butterfly"

if forager-energy > (crop-capacity-red * 5.814)	;; 50 mu liter is 290.7 J

[set breed return-stop-foragers-red]

]

ask yellow-foragers [

move-levy-flight

eat-yellow-flower

set flower-stay-time flower-stay-timer

set size 6

set color yellow

set shape "butterfly"

if forager-energy > (crop-capacity-yellow * 5.814)	;; 25 mu liter is 145.35 J

[set breed return-stop-foragers-yellow]

]

ask red-stop-foragers [

set color red

set size 6

set flower-stay-time flower-stay-time - 1

if flower-stay-time < 0

[ ifelse random 100 > within-trip-constancy

[set breed generalist-bees ]

[set breed red-foragers ]

]

]

ask yellow-stop-foragers [

set color yellow

set size 6

set flower-stay-time flower-stay-time - 1

if flower-stay-time < 0

[ ifelse random 100 > within-trip-constancy

[set breed generalist-bees ]

[set breed yellow-foragers ]

]

]

ask return-stop-foragers-red [

set color red

set size 6

set flower-stay-time flower-stay-time - 1

set distance-red distancexy 0 0

if flower-stay-time < 0

[set breed return-red-foragers]

]

ask return-stop-foragers-yellow [

set color yellow

set size 6

set flower-stay-time flower-stay-time - 1

set distance-yellow distancexy 0 0

if flower-stay-time < 0

[set breed return-yellow-foragers]

]

ask return-stop-foragers-generalist [

set color white

set size 6

set flower-stay-time flower-stay-time - 1

;set distance-red distancexy 0 0

if flower-stay-time < 0

[set breed return-generalist-foragers]

]

ask return-red-foragers [

wiggle

move

set size 6

set color red

set shape "butterfly"

set trip-red 1

set inside-nest-stay-time nest-stay-time

return-to-nest

if pcolor = white [

ifelse random-float recruitment-is-1 < recruitment-probability-red

[ set breed red-influencers]

[set breed inside-nest-workers-red ]]]

ask return-yellow-foragers [

wiggle

move

set size 6

set color yellow

set shape "butterfly"

set trip-yellow 1

set inside-nest-stay-time nest-stay-time

return-to-nest

if pcolor = white [

ifelse random-float 0 > recruitment-probability-yellow	;; no recruitment

[ set breed yellow-influencers]

[set breed inside-nest-workers-yellow ]]]

ask return-generalist-foragers [

wiggle

move

set size 6

set color white

set shape "butterfly"

set trip-generalist 1

set inside-nest-stay-time nest-stay-time

return-to-nest

if pcolor = white [

set breed inside-nest-workers-generalist

]]

ask inside-nest-workers-red [

set color pink

move-inside-nest

wiggle

set shape "butterfly"

set size 6

set inside-nest-stay-time inside-nest-stay-time - 1

ifelse inside-nest-stay-time < 0

[set breed red-foragers

if random 100 > probability-of-constancy [ set breed generalist-bees]

reset-times ]

[ unload-red ]]

ask inside-nest-workers-yellow [

set color pink

move-inside-nest

wiggle

set shape "butterfly"

set size 6

set inside-nest-stay-time inside-nest-stay-time - 1

ifelse inside-nest-stay-time < 0

[set breed yellow-foragers

if random 100 > probability-of-constancy [ set breed generalist-bees]

reset-times ]

[ unload-yellow ]]

ask inside-nest-workers-generalist [

set color yellow

move-inside-nest

wiggle

set shape "butterfly"

set size 6

set inside-nest-stay-time inside-nest-stay-time - 1

if inside-nest-stay-time < 0

[set breed generalist-bees

unload-generalist ]]

ask red-influencers [

set color blue

move-inside-nest

target-yellow-bees

wiggle

set shape "butterfly"

set size 6

set inside-nest-stay-time inside-nest-stay-time - 1

ifelse inside-nest-stay-time < 0

[ set breed red-foragers

if random 100 > probability-of-constancy [ set breed generalist-bees]

reset-times ]

[ unload-red ]]

ask yellow-influencers [

set color blue

move-inside-nest

target-red-bees

wiggle

set shape "butterfly"

set size 6

set inside-nest-stay-time inside-nest-stay-time - 1

ifelse inside-nest-stay-time < 0

[ set breed yellow-foragers

if random 100 > probability-of-constancy [ set breed generalist-bees]

reset-times ]

[ unload-yellow ]]

tick

my-update-plots

end

;;;;;;;;;;;;;;;;;;;;;;;;;;;;;;;;;;;;;;;;;;; behaviour programming ;;;;;;;;;;;;;;;;;;;;;;;;;;;;;;;;

to wiggle

right random 50

left random 50

end

to move

forward 1.4

set forager-energy forager-energy - MetabolicCosts

end

to move-levy-flight

set forager-energy forager-energy - MetabolicCosts

if flight-length <= 0 [

rt random 360

set flight-length levy levy-mu

]

ifelse flight-length < 1.4 [

fd flight-length

set flight-length 0

][

fd 1.4

set flight-length flight-length - 1.4

]

End

to-report levy [ p-mu ]

let result 0

ifelse p-mu <= 1 [

set result random-float 100000000000

][

ifelse p-mu >= 3 [

set result abs random-normal 5 1;0 1

][

let a random-float 1

set result exp ( ln (a) * (1 / 1 - p-mu))

]

]

report result

end

to record-patch-distance

if ticks > ticks-burn-in [

set foraging-distance foraging-distance + patch-distance

set patch-distance 0]

end

to record-patch-distance-red

if ticks > ticks-burn-in [

set foraging-distance-red foraging-distance-red + patch-distance-red

set patch-distance-red 0]

end

to record-patch-distance-yellow

if ticks > ticks-burn-in [

set foraging-distance-yellow foraging-distance-yellow + patch-distance-yellow

set patch-distance-yellow 0]

end

to record-flower-visit

if ticks > ticks-burn-in [

set total-visits total-visits + flower-visits

set flower-visits 0]

end

to record-flower-visit-red

if ticks > ticks-burn-in [

set total-visits-red total-visits-red + flower-visits-red

set flower-visits-red 0]

end

to record-flower-visit-yellow

if ticks > ticks-burn-in [

set total-visits-yellow total-visits-yellow + flower-visits-yellow

set flower-visits-yellow 0]

end

to eat-red-flower

if any? red-flowers-here [

set forager-energy forager-energy + (reward-size-red-flowers * 5.814)

set breed red-stop-foragers

set patches-visited-per-trip patches-visited-per-trip + 1

]

end

to eat-yellow-flower

if any? yellow-flowers-here [

set forager-energy forager-energy + (reward-size-yellow-flowers * 5.814)

set breed yellow-stop-foragers

set patches-visited-per-trip patches-visited-per-trip + 1

]

end

to unload-red	;; procedure at the nest: transfer forager energy to nest

set nest-energy nest-energy + forager-energy

set forager-energy 0 ;; reset forager-energy to 0 (all energy has been transferred to nest)

set foraging-trips-red-bees foraging-trips-red-bees + trip-red

set trip-red 0

set total-patches total-patches + patches-visited-per-trip

set patches-visited-per-trip 0

end

to unload-yellow	;; procedure at the nest: transfer forager energy to nest

set nest-energy nest-energy + forager-energy

set forager-energy 0 ;; reset forager-energy to 0 (all energy has been transferred to nest)

set foraging-trips-yellow-bees foraging-trips-yellow-bees + trip-yellow

set trip-yellow 0

set total-patches total-patches + patches-visited-per-trip

set patches-visited-per-trip 0

end

to unload-generalist	;; procedure at the nest: transfer forager energy to nest

set nest-energy nest-energy + forager-energy

set forager-energy 0 ;; reset forager-energy to 0 (all energy has been transferred to nest)

set foraging-trips-generalist-bees foraging-trips-generalist-bees + trip-generalist

set trip-generalist 0

set total-patches total-patches + patches-visited-per-trip

set patches-visited-per-trip 0

end

to move-inside-nest

forward 0.1

if pcolor = green [rt 180 ]

;set forager-energy forager-energy

end

to target-yellow-bees

if any? inside-nest-workers-yellow-here [

let target one-of inside-nest-workers-yellow-here

ask target [ set breed red-foragers ]

]

end

to target-red-bees

if any? inside-nest-workers-red-here [

let target one-of inside-nest-workers-red-here

ask target [ set breed yellow-foragers ]

]

end

to be-eaten-red

set flower-visits 1

set flower-visits-red 1

if any? generalist-bees-here [

set breed empty-red-flowers

]

if any? red-foragers-here [

set breed empty-red-flowers

]

end

to be-eaten-yellow

set flower-visits 1

set flower-visits-yellow 1

if any? generalist-bees-here [

set breed empty-yellow-flowers

]

if any? yellow-foragers-here [

set breed empty-yellow-flowers

]

end

to return-to-nest

let scent-ahead nest-scent-at-angle	0

let scent-right nest-scent-at-angle 45

let scent-left nest-scent-at-angle -45

if (scent-right > scent-ahead) or (scent-left > scent-ahead)

[ ifelse scent-right > scent-left

[ rt 45 ]

[ lt 45 ] ]

end

to-report nest-scent-at-angle [angle]

let p patch-right-and-ahead angle 1

if p = nobody [ report 0 ]

report [nest-scent] of p

end

to-report recruitment-probability-red

let z exp (logisticA + (logisticB * distance-red))

let recruit-prob z / (1 + z)

report recruit-prob

end

to-report recruitment-probability-yellow

let z exp (logisticA + (logisticB * distance-yellow))

let recruit-prob z / (1 + z)

report recruit-prob

end

to reset-times

set inside-nest-stay-time nest-stay-time

set empty-flower-time flower-refilling-time

end

to my-update-plots

set-current-plot "population"

set-current-plot-pen "red-foragers"

plot count red-foragers

set-current-plot-pen "yellow-foragers"

plot count yellow-foragers

set-current-plot-pen "generalist-bees"

plot count generalist-bees

end
